# Hyperactive WNT/CTNNB1 signaling induces a competing cell proliferation and epidermal differentiation response in the mouse mammary epithelium

**DOI:** 10.1101/2021.06.22.449461

**Authors:** Larissa Mourao, Amber L. Zeeman, Katrin E. Wiese, Anika Bongaarts, Lieve L. Oudejans, Isabel Mora Martinez, Yorick B.C. van de Grift, Jos Jonkers, Renée van Amerongen

## Abstract

In the past forty years, the WNT/CTNNB1 signaling pathway has emerged as a key player in mammary gland development and homeostasis. While also evidently involved in breast cancer, much unclarity continues to surround its precise role in mammary tumor formation and progression. This is largely due to the fact that the specific and direct effects of hyperactive WNT/CTNNB1 signaling on the mammary epithelium remain unknown. Here we use a primary mouse mammary organoid culture system to close this fundamental knowledge gap. We show that hyperactive WNT/CTNNB1 signaling induces competing cell proliferation and differentiation responses. While proliferation is dominant at lower levels of WNT/CTNNB1 signaling activity, higher levels cause reprogramming towards an epidermal cell fate. We show that this involves *de novo* activation of the epidermal differentiation cluster (EDC) locus and we identify master regulatory transcription factors that likely control the process. This is the first time that the molecular and cellular dose-response effects of WNT/CTNNB1 signaling in the mammary epithelium have been dissected in such detail. Our analyses reveal that the mammary epithelium is exquisitely sensitive to small changes in WNT/CTNNB1 signaling and offer a mechanistic explanation for the squamous differentiation that is observed in some WNT/CTNNB1 driven tumors.

## INTRODUCTION

The mammary gland is a defining feature of all mammalian species. It is a dynamic tissue that largely develops after birth, undergoing major changes in proliferation and differentiation during puberty, pregnancy and, ultimately, lactation, when it supplies milk to the newborn offspring. While largely driven by systemic hormones, local signaling cues are equally important to control cell division and cell fate choices during tissue remodeling at each of these stages. Among these is the WNT/CTNNB1 signal transduction pathway, which plays an essential role throughout development of the tissue, from the earliest initiation of mammary placode formation (Chu et al., 2004; Veltmaat et al., 2004) to adult mammary stem cell maintenance (Zeng and Nusse, 2010) and the promotion of side-branching in early pregnancy (Brisken et al., 2000).

Hyperactivation of the WNT/CTNNB1 pathway promotes mammary tumor formation in mice (Nusse and Varmus, 1982). Some, but not all of these tumors display features of squamous differentiation (Miyoshi et al., 2002). How this characteristic is related to WNT/CTNNB1 activity remains unknown due to the inherent latency of tumor development. The involvement of WNT/CTNNB1 signaling in human breast cancer remains less clear (van Schie and van Amerongen, 2020). In general, the direct and immediate effects of elevated WNT/CTNNB1 signaling on the mammary epithelium are poorly understood.

Here we take advantage of a primary organoid culture system that supports short-term growth of the mammary epithelium in the absence of any exogenous growth factors, allowing us to modulate activity of the WNT/CTNNB1 pathway and measure the consequences at a molecular and cellular level. These experiments reveal that WNT/CTNNB1 signaling operates within a precise and narrow range to offer a proliferative advantage to mammary epithelial cells, while higher levels rapidly induce transdifferentiation of the mammary epithelium towards an epidermal cell fate.

## RESULTS

### WNT/CTNNB1 signaling induces size and shape changes in the mammary epithelium

The adult virgin mouse mammary gland is composed of a ductal epithelial network immersed within a stromal fat pad that is mainly composed of adipocytes. The bilayered ductal epithelium contains an outer layer of differentiated myoepithelial and less differentiated progenitor cells (jointly referred to as basal cells) and an inner layer of mature luminal epithelial cells and luminal progenitors. The two cell layers, which originate from a common embryonic progenitor, can be discriminated by the expression of specific cytokeratins, with basal cells expressing KRT5 and KRT14 and luminal cells expressing KRT8 (Figure 1A). Short-term primary organoid cultures of epithelial fragments grown in a basement membrane matrix (Matrigel) and minimal media (Ewald et al., 2008) offer an accessible 3D *in vitro* system for investigating how specific signaling cues affect growth of the mammary gland parenchyma. This is especially relevant for studying the response to changes in WNT/CTNNB1 signaling, since longer term cultures aimed at maintaining mammary epithelial stem cells require WNT/CTNNB1 pathway activation (Jardé et al., 2016; Sachs et al., 2018).

**Figure 1.**
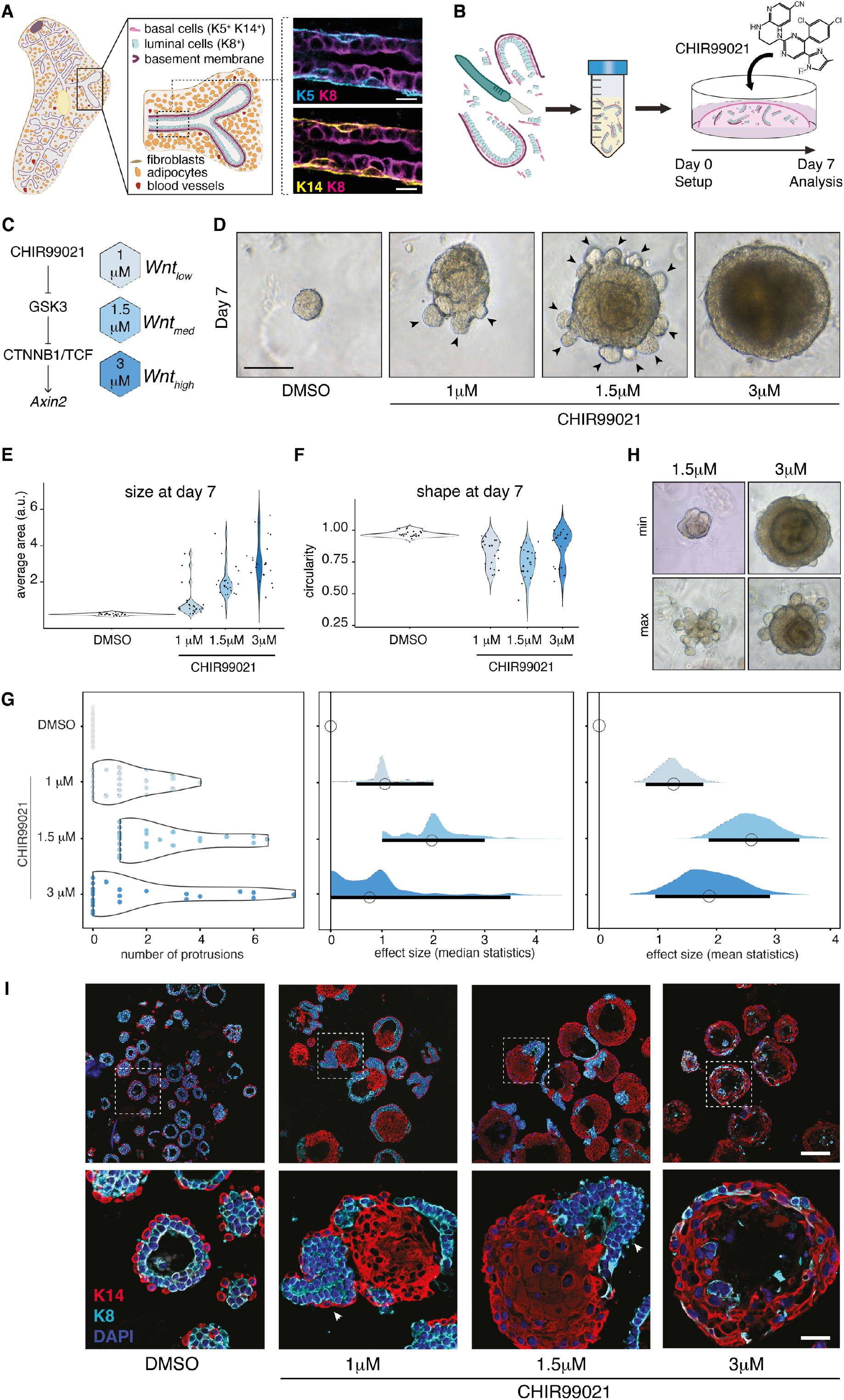
Hyperactivation of the WNT/CTNNB1 pathway induces shape and size changes in the mouse mammary epithelium. A) Cartoon illustrating the cellular composition of the mouse mammary gland and close up of an immunofluorescent staining of the ductal epithelium, depicting the basal (KRT5+ or KRT14+) and luminal (KRT8+) cell layers. Scalebar = 10 μm. B) Schematic illustrating the experimental set up of the 3D primary organoid culture system in which epithelial fragments are embedded in growth-factor reduced matrigel and grown in minimal media for 7 days. C) Addition of the small molecule GSK3 inhibitor CHIR99021 allows dose-dependent hyperactivation of WNT/CTNNB1 signaling, resulting in WNT_low_ (1 μM), WNT_med_ (1.5 μM) and WNT_high_ (3 μM) conditions. D) Representative brightfield microscopy image illustrating the organoid phenotype after 7 days of control (DMSO) or CHIR99021 treatment. Arrowheads point at protrusions. Scalebar = 100 μm. E-F) Violin plots depicting E) the increase in size and F) the change in shape as measured on brightfield microscopy images taken on day 7. Data from n=22 independent organoid cultures are plotted. G) Plots of differences showing (left) the number of protrusions observed (each data point represents the median number of protrusions for one of n=22 independent organoid cultures) and the calculated effect sizes using (middle) median and (right) mean statistics. H) Brightfield microscopy image illustrating the variation in the phenotypes observed in independent organoid cultures. Min = minimal protrusion formation. Max = maximal protrusion formation. I) Confocal microscopy image of an immunofluorescent staining of formalin fixed, paraffin embedded sections of agarose-mounted organoid cultures on day 7. K14 = KRT14 (basal marker), K8 = KRT8 (luminal marker), DAPI = nuclei. Arrowheads point at protrusions. Scalebar = 100 μm (overview) and 25 μm (inserts). Source data and statistics provided in Supplementary File 3.

We followed established protocols (Nguyen-Ngoc et al., 2015), in which freshly digested mammary epithelial fragments are embedded in growth-factor reduced matrigel and cultured in minimal media devoid of WNT signals for 7 days (Figure 1B). In this controlled setup, we induced a dose-dependent increase in WNT/CTNNB1 signaling using different concentrations of CHIR99021, a specific small molecule GSK3 inhibitor that potently activates the WNT/CTNNB1 pathway downstream of the WNT/FZD receptor complex (Figure 1C) (Naujok et al., 2014). While mammary organoids grown under control conditions did not change morphology over the course of the experiment, organoids cultured with increasing concentrations of CHIR99021 (1μM, 1.5μM and 3μM representing Wnt_low_, Wnt_med_ and Wnt_high_ conditions) rapidly increased in size (Figure 1D, Supplementary Figure 1A). After 7 days, they were larger (Figure 1E, increase in Wnt_low_ vs control: 3.6-fold, P=4.09E-2; Wnt_med_ vs control: 7.3-fold, P=1.71E-8; Wnt_high_ vs control: 11.9-fold, P=2.51E-18), less circular (Figure 1F, control: mean 0.696, 95% CI 0.96-0.979; Wnt_low_: mean 0.825, 95% CI 0.778-0.872; Wnt_med_: mean 0.741, 95% CI 0.691-0.791; Wnt_high_: mean 0.878, 95% CI 0.818-0.937) and frequently contained one or more rounded protrusions that appeared to emerge from the core structure (Figure 1D). The latter was more prominent in organoids with intermediate levels of WNT/CTNNB1 signaling (Figure 1G), as also reflected by their lower circularity. Considerable variation existed between independent experiments (n=22 different mice) and individual organoids (Figure 1H, Supplementary Figure 1B, Supplementary File 1).

To characterize the observed phenotypes in more detail, we performed immunofluorescence staining on paraffin embedded organoids. We first probed the expression of two common basal and luminal cell markers, K14 and K8, respectively. Vehicle-treated organoids were organized in two distinct and well-organized cell compartments, with an outer layer of K14^+^ basal cells and an inner cell population of K8^+^ luminal cells as previously described (Ewald et al., 2008). Most organoids also had a defined lumen (Figure 1I). In contrast, organoids with hyperactive WNT/CTNNB1 signaling lost this characteristic epithelial organization. After 7 days, CHIR99021-treated organoids showed separation of the K8^+^ and K14^+^ cell populations, with the K8^+^ cells clustering on the outside, corresponding to the regions where we observed the protrusions (Figure 1I, Supplementary Figure 2A). This is reminiscent of tissue separation via differential adhesion (Foty and Steinberg, 2013). Especially at higher levels of WNT/CTNNB1 signaling, individual nuclei also lost their original shape, size and orientation (Supplementary Figure 2B).

Our combined observations so far suggest that the mouse mammary epithelium shows a complex response to hyperactivation of the WNT/CTNNB1 pathway: The increase in size suggests a proliferative response, while the morphological rearrangement of the basal and luminal compartments suggests a change in cell identity. Of note, these changes occur within a narrow dose-response window, since in our experimental system the difference between the Wnt_high_ (3μM CHIR99021) and Wnt_low_ (1μM CHIR99021) conditions is only three-fold.

### Elevated levels of WNT/CTNNB1 signaling promote cell proliferation

To obtain an unbiased view of the molecular events underlying the observed cellular changes, we performed bulk RNA sequencing on three independent primary organoid cultures harvested at day 7, each containing all four experimental conditions (control, Wnt_low_, Wnt_med_ and Wnt_high_). Given the variation we observed between experiments (Figure 1), we selected three experiments that covered the full range of phenotypes (Supplementary Figure 1B, Supplementary Figure 3A,B), reasoning that this should allow us to identify the most consistent transcriptomic changes. Differential gene expression analysis revealed major changes between the four experimental treatments (Supplementary File 1). Unsupervised hierarchical clustering suggested distinct thresholds for gene activation and repression depending on the absolute levels of WNT/CTNNB1 signaling (Figure 2A).

**Figure 2.**
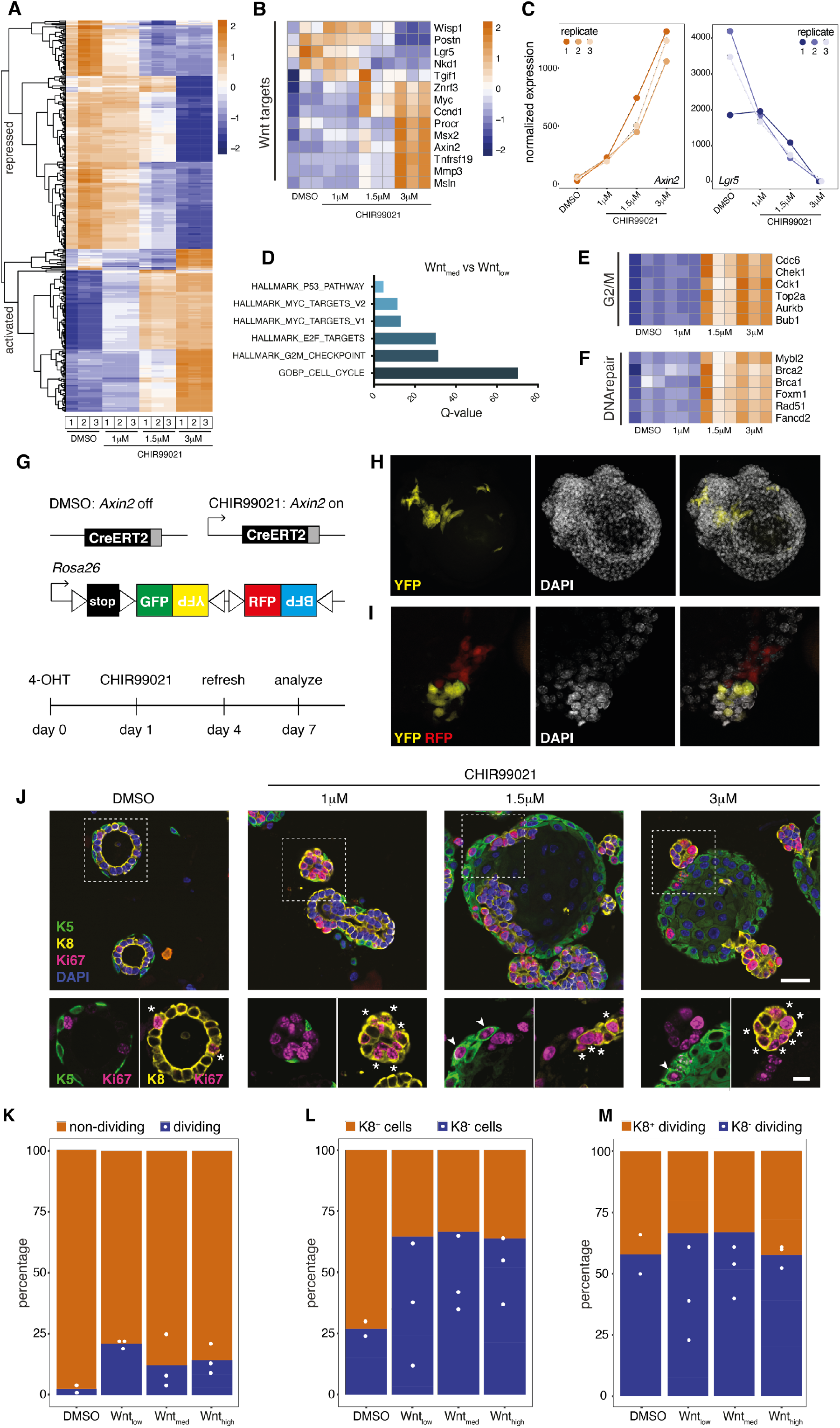
Low levels of WNT/CTNNB1 signaling induce proliferation of basal and luminal cells. A) Heatmap (unsupervised clustering, log2-transformed normalized expression values, Z-score) showing distinct thresholds for gene expression changes at different levels of WNT/CTNNB1 signaling activity. For this and all following heatmaps, RNAseq results for n=3 independent organoid cultures are depicted for all treatment conditions. B) Heatmap (unsupervised clustering, Z-score) of WNT/CTNNB1 target genes that are differentially expressed in one or more conditions. C) Graph showing normalized expression values of the three RNAseq replicates for two of the WNT/CTNNB1 target genes from B): *Axin2* (left) and *Lgr5* (right). D) Bar plot depicting the results of a gene set enrichment analysis for genes that are differentially expressed in WNT_med_ (1.5 μM CHIR99021) versus WNT_low_ (1 μM CHIR99021) organoids. E-F) Heatmaps (unsupervised clustering, Z-score) showing a selection of differentially expressed genes involved in E) the G2/M checkpoint and F) DNA repair. G) Schematic illustrating the lineage tracing setup for the experiments depicted in H-I. A tamoxifen-inducible CreERT2 recombinase is expressed in *Axin2*-positive cells, allowing recombination of a *Rosa26-Confetti* multicolor reporter allele in cells with active WNT/CTNNB1 signaling (i.e. only in the presence of CHIR99021). H-I) Confocal microscopy images showing clonal outgrowth of WNT/CTNNB1-responsive cells using the setup depicted in G. J) Confocal microscopy image of an immunofluorescent staining of formalin fixed, paraffin embedded sections of agarose-mounted organoid cultures on day 7. K5 = KRT5 (basal marker), K8 = KRT8 (luminal marker), KI67 = cell proliferation marker, DAPI = nuclei. Asterisks point to dividing luminal cells. Arrowheads point to dividing basal cells. Scalebar = 25 μm (overview) and 10 μm (inserts). K-M) Stacked bar graphs showing quantification of n=2 (DMSO) and n=3 (CHIR) independent experiments similar to the one depicted in J. Total cell numbers counted per experiment (regions of interest (ROI) based on DAPI signal): DMSO: 655 and 1957; WNT_low_: 529, 9473 and 1261; WNT_med_: 6294, 240 and 1675; WNT_high_: 1942, 478 and 398. Source data and statistics provided in Supplementary File 3.

We first confirmed that WNT/CTNNB1 signaling is indeed hyperactivated in CHIR99021 treated organoids. To this end, we analyzed the response pattern of a curated set of 21 genes that have previously been reported to be activated by WNT/CTNNB1 signaling in the mammary gland or in other tissues (Boonekamp et al., 2021; Fafilek et al., 2013; Szemes et al., 2018; Wang et al., 2015; Yu et al., 2016). We made sure to include multiple bona-fide feedback targets, including *Axin2* and *Lgr5* (Supplementary File 2). Out of these 21 genes, 14 were differentially expressed in one or more conditions (Figure 2B). Most showed the expected dose-dependent induction, including *Axin2* and the cell proliferation markers *Ccnd1* and *Myc*, although the absolute levels and dynamic range varied between genes (Supplementary File 3). One notable exception was *Lgr5*, the expression of which was consistently downregulated in all replicates (average 790-fold change in Wnt_high_ versus control, Figure 2C).

Compared to Wnt_low_ organoids, Wnt_med_ organoids were enriched for genes involved in cell cycle progression (Figure 2D, Supplementary File 2, p-value = 1.43E-75, FDR q-value = 4.61E-71), most notably the G2/M checkpoint (p-value = 1.07E-33, FDR q-value = 5.35E-32). Almost 90 differentially expressed genes were classified as targets of the dimerization partner, RB-like, E2F and multi-vulval class B (DREAM) complex, which regulates cell-cycle dependent gene expression (Sadasivam and DeCaprio, 2013) (Supplementary File 2, p-value = 1.43e-75, FDR q-value = 4.61e-71). This includes multiple E2F4 targets, such as *Top2a* and *Aurkb*, suggesting that their repression is relieved in response to enhanced WNT/CTNNB1 signaling (Figure 2E). Of note, the comparison of Wnt_low_ and Wnt_med_ expression profiles also revealed the induction of homologous recombination and Fanconi anemia pathway genes, such as *Rad51*, *Brca1* and *Fancd2* (Figure 2F), which were recently shown to be induced in WNT addicted cancers in a CTNNB1-and MYBL2/FOXM1 dependent manner (Kaur et al., 2021). Indeed, both *Mybl2* and *FoxM1* were upregulated in Wnt_med_ organoids as well (Figure 2F), suggesting that the same mechanism might operate in direct response to elevated levels of WNT/CTNNB1 signaling.

To confirm that the mammary epithelial cells indeed divide as a result of increased WNT/CTNNB1 signaling, we performed lineage tracing in primary organoids derived from *Axin2^CreERT2^;Rosa26^Confetti^* mice (Van Amerongen et al., 2012; Snippert et al., 2010), in which Cre/lox mediated recombination of a multi-color fluorescent reporter allele can be induced in cells that express the WNT/CTNNB1 target gene *Axin2* (Figure 2G). In the combined presence of CHIR99021 (which hyperactivates WNT/CTNNB1 signaling) and 4-hydroxytamoxifen (which induces Cre^ERT2^ recombinase activity) clonal cell proliferation occurred in both basal and luminal regions, with large, flattened cells accumulating in the center of the organoid (Figure 2H-I, Supplementary Figure 2C).

To validate our findings, we performed staining for the cell proliferation marker Ki67. This showed that, as expected, very few cells were dividing in control treated organoids, with 2% of all luminal cells and 6% of basal cells dividing (Supplementary File 3). Both dividing basal and luminal cells could be detected in organoids with hyperactive WNT/CTNNB1 signaling, but they were largely absent from the center (Figure 2J). Quantification of the number of dividing luminal and non-luminal cells across the different conditions revealed the highest proportion of dividing cells in Wnt_low_ organoids (mean 21% versus mean 2.5% in DMSO treated cells, P=0.02, Figure 2K), suggesting that the increase in organoid size (highest in Wnt_high_, Figure 1F) cannot solely be due to an increase in cell proliferation. The overall proportion of luminal cells showed a slight but steady decrease with higher levels of WNT/CTNNB1 signaling (mean 73% of all cells in DMSO versus mean 48% of all cells in Wnt_high_, Figure 2L), but we did not detect statistically significant changes in the division of luminal and non-luminal cells across the different treatment conditions (Figure 2M and Supplementary File 3). Thus, we conclude that increased levels of WNT/CTNNB1 signaling promote cell division in both basal and luminal mammary epithelial cells.

### Hyperactive WNT/CTNNB1 signaling induces squamous differentiation

Returning to our transcriptomics analysis, we next asked if we could identify specific gene signatures with distinct dose-dependent response patterns. To this end, we performed fuzzy c-means clustering (Kumar and Futschik, 2007) on all genes that were differentially expressed in one or more conditions. This allowed us to identify 12 clusters with co-regulated genes (Supplementary Figure 3C,D, Supplementary File 2). Functional annotation and gene set enrichment analysis of individual clusters revealed the dose-dependent and counterintuitive loss of a mammary stem cell signature (cluster 2, Figure 3A, Supplementary File 2, LIM_MAMMARY_STEM_CELL_UP: p-value = 2.05e-14, FDR q-value = 5.02e-10) (Lim et al., 2010) and the loss of contractile features (cluster 10, Figure 3B, Supplementary File 2, GOCC_CONTRACTILE_FIBER: p-value = 1.07e-26, FDR q-value = 1.73e-22). This included the loss of myoepithelial cell markers *Acta2* and *Otxr* (Figure 3C, average fold reduction of 27.6 and 160 respectively in Wnt_high_ vs control, Supplementary File 3). The expression of basal keratins *Krt14* and *Krt5* increased rather than decreased (Figure 3D, average fold increase of 12.1 and 9 respectively in Wnt_high_ vs control, Supplementary File 3). Although this may partially be due to the loss of luminal cells, this suggests that basal cells specifically lose their myoepithelial fate, but remain present as such, as also supported by our immunostaining experiments (Figure 1I).

**Figure 3.**
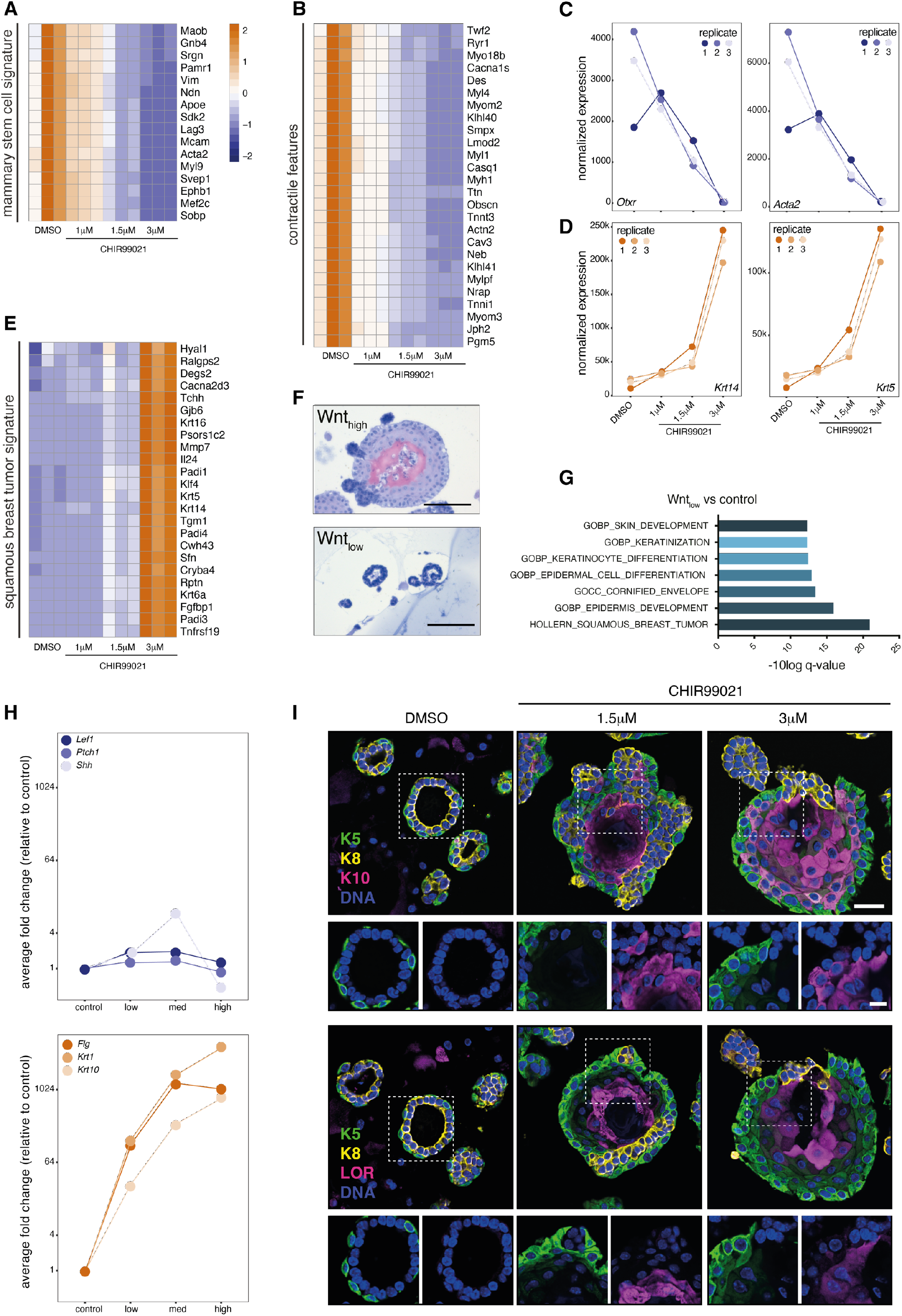
High levels of WNT/CTNNB1 signaling induce squamous differentiation. A-B) Heatmaps (unsupervised clustering, Z-score) showing gradual loss of A) a mammary stem cell signature and B) myoepithelial features with increasing levels of WNT/CTNNB1 signaling. C-D) Graphs showing normalized expression values of the three RNAseq replicates for C) two myoepithelial markers (*Otxr* and *Acta2*) and D) two basal markers (*Krt14* and *Krt5*). E) Heatmap (unsupervised clustering, Z-score) showing gain of a squamous breast tumor signature with increasing levels of WNT/CTNNB1 signaling. F) Brightfield microscopy image showing H&E staining of paraffin embedded organoids, which reveals a core of keratinized material in the center of WNT_high_ organoids. G) Bar plot depicting the results of a gene set enrichment analysis for genes that are differentially expressed in WNT_low_ (1 μM CHIR99021) versus control (DMSO) organoids. H) Graphs showing the average fold change in hair matrix (top: *Lef1*, *Ptch1*, *Shh*) and skin barrier markers (bottom: *Flg*, *Krt1*, *Krt10*). Data points depict the mean of the three RNAseq replicates for each experimental condition, with the control (DMSO) set to 1. I) Confocal microscopy image of an immunofluorescent staining of formalin fixed, paraffin embedded sections of agarose-mounted organoid cultures on day 7. K5=KRT5 (basal marker), K8 = KRT8 (luminal marker), K10 = KRT10 (suprabasal marker), LOR = Loricrin (terminal differentiation marker), DAPI = nuclei. Scalebar = 25 μm (overview) and 10 μm (inserts). Source data provided in Supplementary File 3.

Among the gene clusters that predominantly increased in expression upon hyperactivation of the WNT/CTNNB1 pathway, one stood out by showing hallmarks of squamous differentiation, which included the gain of a squamous breast tumor signature (cluster 3, Figure 3E, Supplementary File 2, HOLLERN_SQUAMOUS_BREAST_TUMOR: p-value = 4.8e-31, FDR q-value = 7.75e-27). Closer inspection revealed upregulation of multiple genes associated with epidermal development, keratinization and cornification (cluster 3, Supplementary File 2, GOBP_CORNIFICATION: p-value = 4.79e-13; FDR q-value = 7.03e-10, GOBP_KERATINIZATION: p-value = 5.43e-13; FDR q-value = 7.63e-10, GOBP_EPIDERMIS_DEVELOPMENT: p-value = 2.33e-12; FDR q-value =2.42e-09). Indeed, H&E stained organoids sections revealed a dense core of eosinophilic material in the center of Wnt_high_ but not Wnt_low_ organoids (Figure 3F), resembling keratin pearls that are characteristic of squamous differentiation *in vivo* (Supplementary Figure 2D). While squamification was most apparent in the Wnt_high_ condition, a squamous gene expression signature was also already present in Wnt_low_ organoids (Supplementary File 2, Figure 3G). This suggests that squamous differentiation and proliferation of the mammary epithelium are induced in parallel, with the squamous phenotype becoming more dominant at higher levels of WNT/CTNNB1 signaling.

### Reprogramming of the mouse mammary epithelium towards an epidermal cell fate

The mammary gland develops as a skin appendage. Like the hair follicle, it starts as a local thickening of the surface ectoderm. WNT signaling is required for the initiation of both mammary and hair follicle placode formation (Andl et al., 2002; Chu et al., 2004) and hyperactivation of WNT/CTNNB1 signaling in the epidermis, via the expression of a dominant active form of CTNNB1 under the control of a *Krt14* promoter, is sufficient to drive *de novo* hair follicle formation and gives rise to hair tumors, or pilomatricomas, in older mice (Gat et al., 1998). At the same time, WNT/CTNNB1 signaling also controls growth and maintenance of non-hairy and interfollicular epidermis (Choi et al., 2013; Lim et al., 2013). We therefore asked if the squamous differentiation we observed reflected reprogramming of the mammary epithelium towards either of these cell fates.

Expression of the hair matrix markers *Lef1*, *Shh* and *Ptch1* (Gat et al., 1998) did not change in response to CHIR99021 treatment, but we did detect increased expression of genes encoding epidermal keratins (*Krt1* and *Krt10*) and skin barrier proteins (*Flg*) (Figure 3H, Supplementary File 3). Immunofluorescence staining confirmed the presence of KRT10-positive cells immediately adjacent to KRT5-positive basal cells as well as the presence of Loricrin (LOR) positive cells in the center of both WNT_med_ and WNT_high_ organoids (Figure 3I). This pattern of expression is identical to that observed in stratified epithelia, where KRT10 is expressed in the first suprabasal (or spinous) layer and where LOR expression switches on in the upper spinous and lower granular layer. These differentiating cells no longer divide, thereby also explaining the lack of proliferation in this area (compare Figure 2J and 3I, (Supplementary Figure 2D). Together, these findings suggest that supraphysiological levels of WNT/CTNNB1 signaling induce transdifferentiation of the mammary epithelium towards an epidermal cell fate.

To find further support for this hypothesis, we took advantage of the existence of scRNAseq gene expression signatures that distinguish different parts of the epidermis and anagen hair follicle (Joost et al., 2020). We investigated the (changes in) expression of the top 20 genes (Supplementary File 2) that characterize 33 distinct subpopulations in either the permanent epidermis (9/33 clusters) or the anagen hair follicle (24/33 clusters). CHIR99021 treated mammary organoids specifically induce markers that characterize the (cycling) basal and suprabasal interfollicular epidermis and the suprabasal upper hair follicle, but not of the sebaceous gland, outer bulge or hair germ (Supplementary Figure 4A-I). CHIR99021 treated organoids also expressed some markers that characterize different parts of the cycling anagen hair follicle (Supplementary Figure 4J-AG). These were mostly associated with actively dividing populations, however (Supplementary Figure 4P-S). Their expression (e.g. *Top2a*, *Ccnb1*, *Cenpf*) therefore likely reflects the increased cell division we observe, rather than a distinct cell fate.

Closer inspection of the observed gene expression changes revealed that in addition to *Krt1* and *Krt10*, the highest increase in keratin gene expression was observed in a select number of keratin genes (*Krt6a*, *Krt6b*, *Krt16*, *Krt17*, Supplementary File 3) that are typically induced in interfollicular epidermis that is stressed or wounded (Zhang et al., 2019). No such increase was observed for keratin genes that characterize other stratified epithelia, such as the palmoplantar epidermis (*Krt9*)(Schweizer et al., 1989) or the oral and esophageal epithelium (*Krt4*, *Krt13*)(van Muijen et al., 1986; Trisno et al., 2018). While analyzing our RNAseq data, we realized that many of the genes associated with the epidermal, keratinization and cornification gene signatures (i.e. cluster 3) were located in the same region, namely the epidermal differentiation complex (EDC) locus on mouse chromosome 3q (Figure 4A). Spanning more than 3Mb, the EDC harbors ∼60 genes that play a critical role in terminal differentiation of the epidermis. It contains 4 distinct gene families, with the *S100* genes found at the 5’ and 3’ border and the small proline-rich (*Sprr*) genes, late cornified envelope (*Lce*) genes and *Flg*-like genes (e.g. *Flg*, *Rptn* and *Tchh*) located in between. While the most proximal and distal *S100* genes (*S100a1, S100a13*, *S100a11* and *S100a10*) were expressed in control treated organoids and remained expressed at comparable levels irrespective of CHIR99021 treatment (Figure 4A,B), expression of the intervening genes was low to undetectable in DMSO treated organoids. Upon hyperactivation of the WNT/CTNNB1 pathway, however, 55 genes belonging to different families (e.g. *Sprr1a*, *Lce1b* and *Rptn*) dramatically increased in expression (Figure 4A,C). This suggests that the entire EDC becomes activated, as would be expected for epidermal keratinocytes. Taking everything together, we conclude that hyperactive WNT/CTNNB1 signaling in primary mammary organoids is sufficient to reprogram cells to an epidermal state, after which the reprogrammed cells undergo the normal differentiation program of basal keratinocytes.

**Figure 4.**
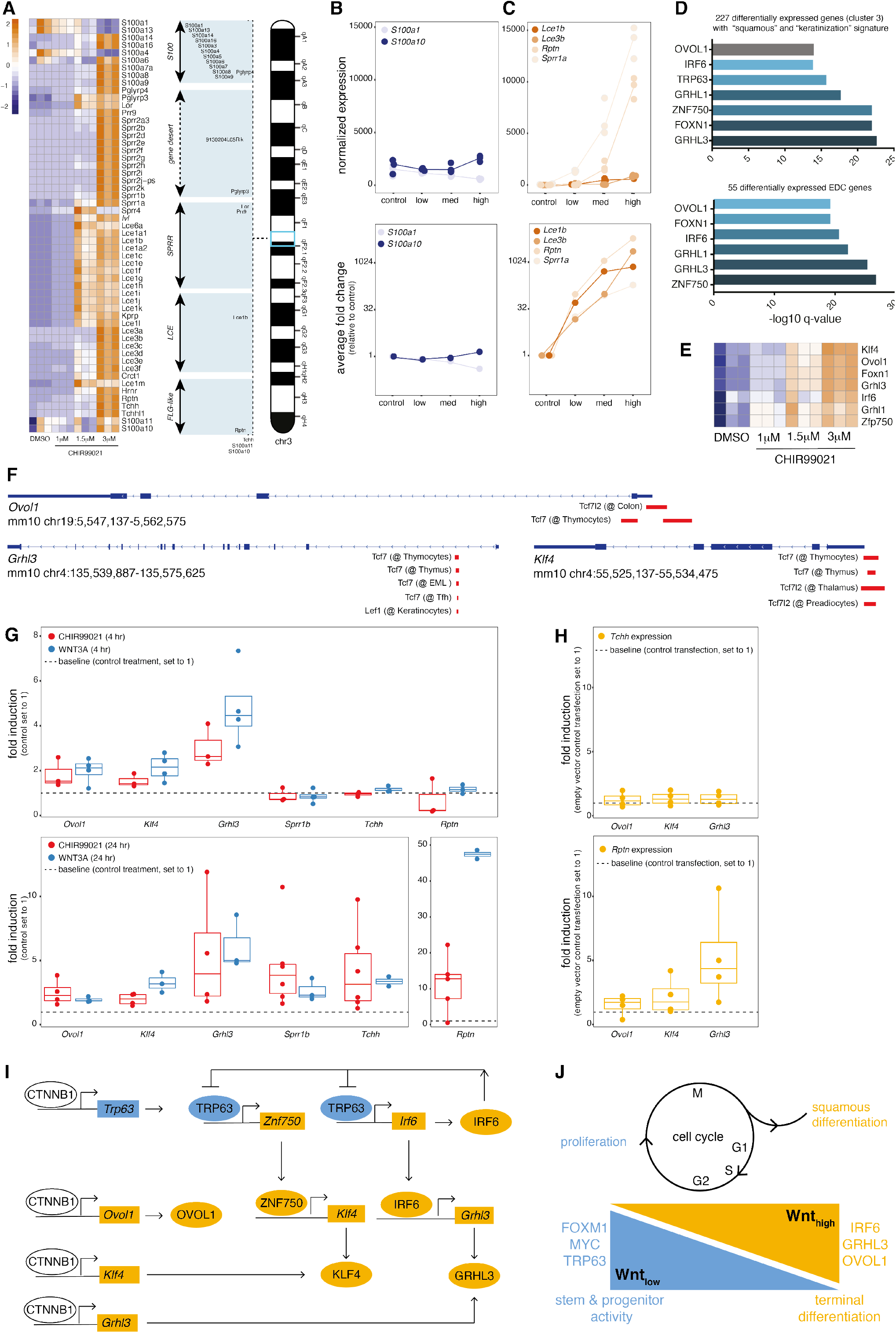
WNT/CTNNB1 signaling induces master regulators of epidermal differentiation. A) Heatmap (genes depicted in order of chromosomal location, Z-score) showing *de novo* expression of multiple EDC locus genes. B-C) Graphs showing normalized expression (top) and average fold change (bottom) of two genes located at the border (*S100a1* and *S100a10*) and four genes from the central region of the EDC locus (*Lce1b*, *Lce3b*, *Rptn*, *Sprr1a*). For normalized expression values individual data points of the three RNAseq replicates are shown. For average fold change values data points depict the mean of the three RNAseq replicates for each experimental condition. D) Bar plot depicting the results of a gene set enrichment analysis for the 227 differentially expressed from cluster 3, which revealed the keratinization signature (top) and for the 55 differentially expressed genes located in the EDC locus (bottom). E) Heatmap (unsupervised clustering, log-transformed normalized expression values, Z-score) of 7 differentially expressed candidate master regulatory transcription factors. F) Schematic depicting the results from ChIPseq analyses, revealing the presence of common TCF/LEF binding sites (red blocks) close to the transcriptional start site of *Ovol1* (top), *Grhl3* (bottom left) and *Klf4* (bottom right). G) Graphs depicting the results of quantitative RT-PCR analyses performed in BC44 cells, revealing the induction of master regulators (*Ovol1*, *Klf4* and *Grhl3*) and a selection of EDC locus genes (*Sprr1b*, *Tchh* and *Rptn*) by both CHIR99021 and purified WNT3A. Data points depict n=2-6 independent experiments, with each data point representing the average of a technical triplicate. *Rpl13a* was used as a reference gene and all expression levels are plotted as fold increase over control (DMSO or BSA treated cells). H) Graphs depicting the results of quantitative RT-PCR analyses performed in BC44 cells, revealing the induction of *Rptn* (bottom) but not *Tchh* (top) following transient overexpression of GRHL3 and, to a lesser extent, OVOL1 and KLF4. Grhl3 and, to a lesser extent, Ovol1 and Klf4. Data points depict n=4 independent experiments, with each data point representing the average of a technical triplicate. *Rpl13a* was used as a reference gene and all expression levels are plotted as fold increase over control (empty vector control transfected cells). I) Model of the gene regulatory network that controls epidermal transdifferentiation. J) Model summarizing the WNT/CTNNB1-induced competing proliferation and differentiation responses of the mammary epithelium. See text for details.

### WNT/CTNNB1 signaling induces master regulators of keratinocyte differentiation

So how does WNT/CTNNB1 signaling transform mammary epithelial cells towards an epidermal fate? We reasoned that this massive change in gene expression likely requires the activity of one or more master regulators of epidermal differentiation. Using gene set enrichment analysis for the 227 differentially expressed genes from cluster 3, which gave rise to the squamous gene signature (Supplementary File 2, Supplementary Figure 3), as well as the 55 differentially expressed EDC genes (Figure 4A), we generated a shortlist of candidate transcription factors (Supplementary File 3). Of the top-ranked candidates, 6 were shared between the two queried gene sets (Figure 4D). All 6 were dose-dependently induced in CHIR99021 treated mammary organoids (Figure 4E). We selected two of these genes (*Ovol1* and *Grhl3*, typically expressed in basal and suprabasal keratinocytes, respectively), together with *Klf4* for experimental follow-up based on their known involvement in skin barrier formation (Segre et al., 1999; Teng et al., 2007; Ting et al., 2005) and their proposed role in EDC locus regulation (Klein et al., 2017; Nascimento et al., 2011). Using published ChIPseq data, we detected multiple common TCF/LEF binding sites close to the transcriptional start site of all three genes (Figure 4F). We therefore speculate that WNT/CTNNB1 signaling directly induces a squamous differentiation program by binding to regulatory elements that control the expression of these master regulatory transcription factors.

To validate that WNT/CTNNB1 signaling indeed induces the expression of *Ovol1*, *Grhl3* and *Klf4* prior to activating EDC locus genes, we stimulated the WNT-responsive BC44 mouse mammary epithelial cell line (Deugnier et al., 1999) with either 3 μM CHIR99021 or 50 ng/ml purified WNT3A protein for 4 or 24 hours. Next, we determined the expression of the three putative master regulators (*Ovol1*, *Klf4* and *Grhl3*) as well as three EDC locus genes (*Sprr1b*, *Tchh* and *Rptn*) by quantitative RT-PCR. After 4 hours, both CHIR99021 and purified WNT3A induced the expression of endogenous *Ovol1*, *Klf4* and *Grhl3* but not endogenous *Sprr1b*, *Rptn* and *Tchh* above baseline (Figure 4G, Supplementary File 3). After 24 hours, the EDC locus genes were also induced, with endogenous *Rptn* showing the highest fold change (47-fold increase in WNT3A treated cells versus control, Figure 4G, Supplementary File 3). Of note, transient overexpression of GRHL3, but not OVOL1 or KLF4, was sufficient to induce endogenous *Rptn* but not its neighboring gene, *Tchh* (Figure 4H, Supplementary File 3). This is in line with the fact that *Rptn* was previously suggested to be a conserved target gene for GRHL transcription factors in both mouse and human (Mathiyalagan et al., 2019).

## DISCUSSION

Using a short-term, 3D primary organoid culture system, we have dissected the early response of the mouse mammary epithelium to elevated levels of WNT/CTNNB1 signaling (Figure 1). We show that hyperactive WNT/CTNNB1 signaling is sufficient to induce proliferation of both basal and luminal cells (Figure 2). At the same time, increased levels of WNT/CTNNB1 signaling induce squamous differentiation (Figure 3). These two activities compete with each other, resulting in specific changes in organoid size, shape and cellular composition that depend on the absolute levels of WNT/CTNNB1 signaling. At lower levels, proliferation has the upper hand, while at higher levels differentiation becomes dominant (Figure 4I). Of note, these events become apparent within 3 days and occur within a narrow dose-response window (Supplementary Figure 1).

The squamous differentiation we observe in our primary organoids resembles the phenotype that was previously reported for a 3D culture system aimed at allowing long-term passaging of mouse mammary organoids. Here, organoids are grown in the presence of RSPO1 (Jardé et al., 2016), suggesting that slight amplification of the endogenous WNT/CTNNB1 signaling activity is already sufficient to transform the mouse mammary epithelium into epidermis. We hypothesize that the consistent and major reduction in *Lgr5* expression levels that we observe in response to WNT/CTNNB1 hyperactivation (Figure 2C) reflects an attempt of the mammary epithelium to bring WNT/CTNNB1 signaling levels back down into to the physiological range. Given that *Lgr5* is typically considered to be a positive WNT/CTNNB1 feedback regulator (Barker et al., 2007), this underscores the dynamic adaptation of such feedback loops.

Our data suggest that the squamous differentiation signature reflects reprogramming of the mammary epithelium towards an epidermal cell fate. This transdifferentiation event involves activation of the EDC locus (Figure 4), which normally only occurs in differentiating keratinocytes and which typically involves physical relocation of the EDC locus from the nuclear periphery to the nuclear interior. This requires major changes in chromatin organization and (super) enhancer activity and is thought to be regulated by the *Trp63*-mediated induction of the SWI/SNF chromatin remodeling factor *Smarca4* (previously called *Brg1*) (Mardaryev et al., 2014), while also possibly involving GRHL3 (Klein et al., 2017; Poterlowicz et al., 2017). Recent work suggests that reduced tension of the nuclear lamina in suprabasal keratinocytes, resulting from the loss of ITGB1 attachment to the extracellular matrix, can also directly induce physical relocation and transcriptional activation of the EDC locus (Carley et al., 2021). Interestingly, we also observe changes in nuclear shape and size in response to hyperactive WNT/CTNNB1 signaling (Supplementary Figure 2). The precise nature of this event remains unknown, but it could well be either a cause or consequence of reprogramming towards and epidermal cell fate.

Furthermore, our data show that WNT/CTNNB1 signaling in and by itself is sufficient to jumpstart a complex gene regulatory network that involves multiple master regulators of epidermal differentiation (Figure 4D-H). Together, these transcription factors are well known to control epidermal differentiation and other ectodermal developmental processes (Figure 4I) (Dai et al., 1998; Ferretti et al., 2011; Kimura-Yoshida et al., 2015; Koster et al., 2004; Nair et al., 2006; Romano et al., 2012). Both *Irf6* and *Znf750* are known TRP63 target genes. They, in turn, are thought to induce the expression of *Klf4* and *Grhl3* as well as terminal differentiation genes (Moretti et al., 2010; Oberbeck et al., 2019; Sen et al., 2012). *Ovol1* and *Trp63* have previously been shown to be directly activated by WNT/CTNNB1 signaling (Ferretti et al., 2011; Li et al., 2002). *Grhl3* has also previously been suggested to be a direct WNT target gene in osteoblasts (Salazar et al., 2016). Our data suggest that a similar gene regulatory network can be induced in mammary epithelial cells.

Of course, many questions remain. First and foremost, what ultimately shifts the balance from proliferation to differentiation remains to be determined (Figure 4J). Mechanistically, the proliferation response is characterized by an expression signature that is enriched for genes involved in the G2/M checkpoint and DNA repair (Figure 2). On the one hand, this could indicate that WNT/CTNNB1 signaling operates to enhance DNA repair, as recently suggested (Kaur et al., 2021). On the other hand, it is tempting to speculate that this signature actually reflects replication stress. This would fit with an earlier observation that increased WNT signaling induces a DNA damage response in primary human mammary epithelial cells (Ayyanan et al., 2006) and could explain why higher levels of WNT/CTNNB1 signaling do not continue to offer a proliferative advantage. This would also fit with the induction of stress keratins (Supplementary File 3). In fact, it might be directly connected to the squamous differentiation phenotype, as keratinocytes are known to undergo differentiation in response to DNA damage and replication stress (Freije et al., 2014; Molinuevo et al., 2020).

One caveat of the current study is that we have not yet resolved the earliest temporal changes in gene expression nor the precise nature of CTNNB1-dependent DNA binding events. Second, it will be interesting to see if a similar response can be detected *in vivo* and, related to this, if basal and luminal cells respond differently. In our experimental setup we cannot clearly discriminate the behavior of basal and luminal cells. While both seem to proliferate in response to hyperactive WNT/CTNNB1 signaling (Figure 2), it is not yet clear if both also undergo squamous differentiation. While basal cells may seem more likely to transdifferentiate, the mammary epithelium develops from a common embryonic progenitor (Spike et al., 2012; Wansbury et al., 2011) and both basal and luminal cells have great inherent plasticity (Van Keymeulen et al., 2015; Koren et al., 2015).

Finally, a logical next question is in how far our findings are relevant for the human mammary epithelium and the development or treatment of breast cancer. In mice, only low to intermediate levels of WNT/CTNNB1 signaling are able to drive mammary tumor formation, with higher levels invariably leading to squamous differentiation and metaplastic tumors (Edwards et al., 1992; Miyoshi et al., 2002; Monteiro et al., 2014; Tsukamoto et al., 1988). In humans, activating genetic mutations resulting in high levels of WNT/CTNNB1 signaling are typically only found in a subset of metaplastic breast carcinomas (Hayes et al., 2008; Ng et al., 2017). Both growth promoting and inhibitory effects of paracrine WNT signaling, including a squamous differentiation response in the latter, have been reported in patient derived xenografts (Green et al., 2013). While squamous differentiation has been reported to occur in human breast cancer (Tsuda et al., 1997), it is not considered to be common. At the same time, mouse squamous tumors and human basal tumors share some gene expression features (Hollern et al., 2018). If and how this correlates to active WNT/CTNNB1 signaling remains unknown. Determining WNT-responsive gene expression signatures for the healthy and cancerous human breast epithelium will be a critical first step to shed more light on this matter. Our data would predict that here too, subtle changes in WNT/CTNNB1 signaling will affect cell proliferation and differentiation in parallel.

In conclusion, our findings highlight the sensitivity of the mammary epithelium to small changes in WNT/CTNNB1 signaling and offer a mechanistic explanation for the selection of ‘just right’ levels of WNT/CTNNB1 signaling in mammary tumor formation (Gaspar et al., 2009; van Schie and van Amerongen, 2020). Accordingly, we hypothesize that human breast tumors will have a Wnt_low_ signature that promotes proliferation. In contrast, cells with a Wnt_high_ signature, which induces reprogramming towards an epidermal phenotype and results in squamous differentiation, are likely to be counter selected at an early stage of tumor development.

## Supporting information

Supplemental File 1 - RNAseq

Supplemental File 2 - gene lists

Supplemental File 3 - source data and statistics

## SUPPLEMENTARY FILES AND SUPPLEMENTARY FIGURE LEGENDS

**Supplementary File 1**

This file contains the output of the RNAseq analysis using DESeq2. It contains lists of differentially expressed genes between the different treatment conditions (6 lists in total, FDR set at <0.1 for each) as well as an overview of the normalized counts for all genes. The raw data for the RNAseq analysis have been deposited in NCBI GEO (GSE178321).

**Supplementary File 2**

This file contains the gene lists that were used for the analyses mentioned in the text, and as depicted in the main figures and Supplementary Figure 4. It also contains the raw results of the gene set enrichment analyses.

**Supplementary File 3**

This file contains all of the source data and statistical analyses that were performed for the analyses mentioned in the text and depicted in the main figures and Supplementary Figure 4.

**Supplementary Figure 1.**
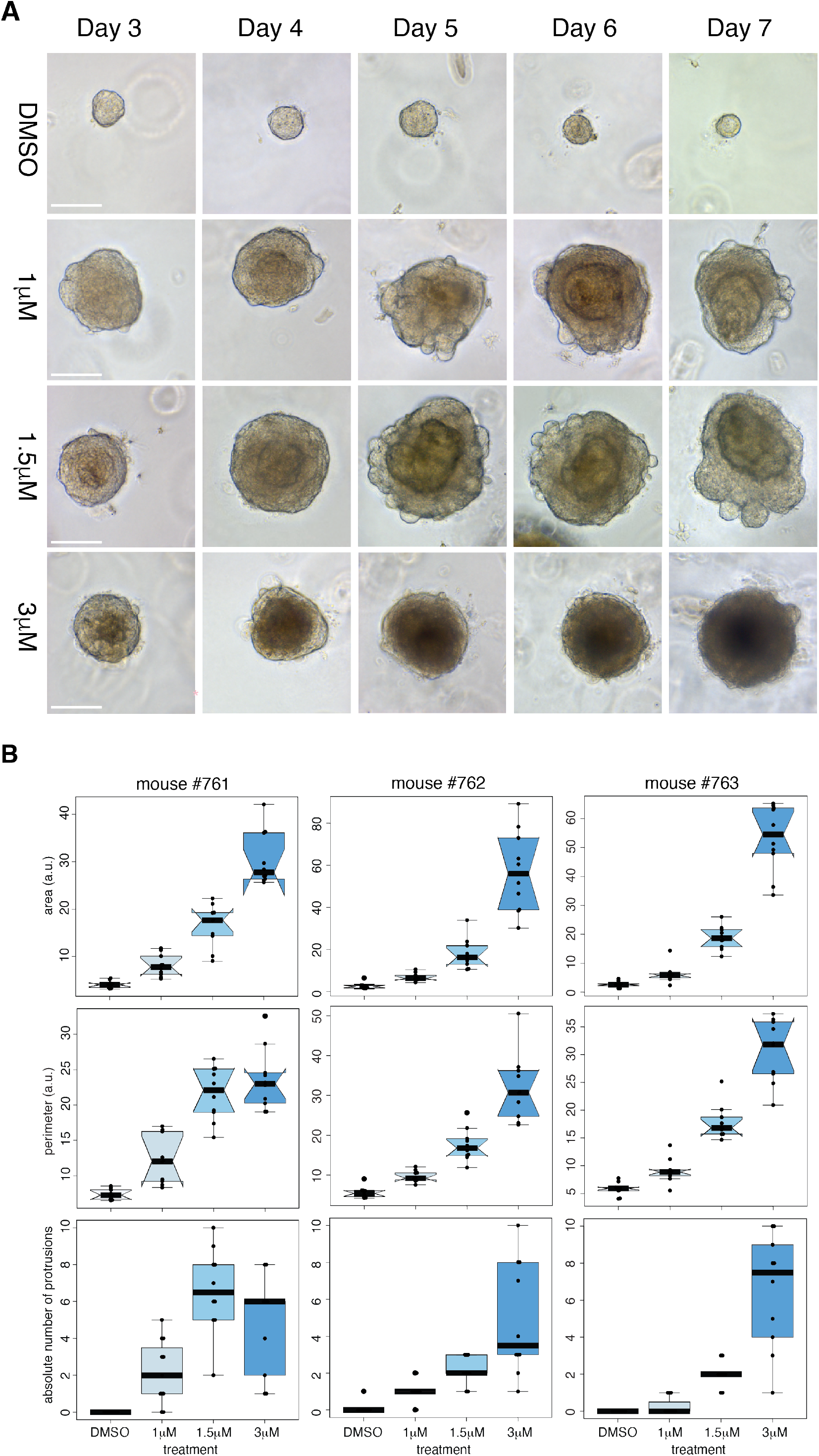
A) Brightfield microscopy images showing temporal changes in size and morphology of a representative organoid culture. Box plots showing changes in size (area, top row, and perimeter, middle row) and morphology (number of protrusions, bottom row) in response to WNT/CTNNB1 hyperactivation. The three samples depicted (mouse #761, #762 and #763) are the samples that were used for the RNAseq experiment. Images from #761 are depicted in 1H as “max” and images from #762 are depicted in 1H as “min”. The RNAseq data deposited at NCBI GEO (GSE178321) are labelled “rep1” (for #763), “rep2” (for #762) and “rep3” (for #761).

**Supplementary Figure 2.**
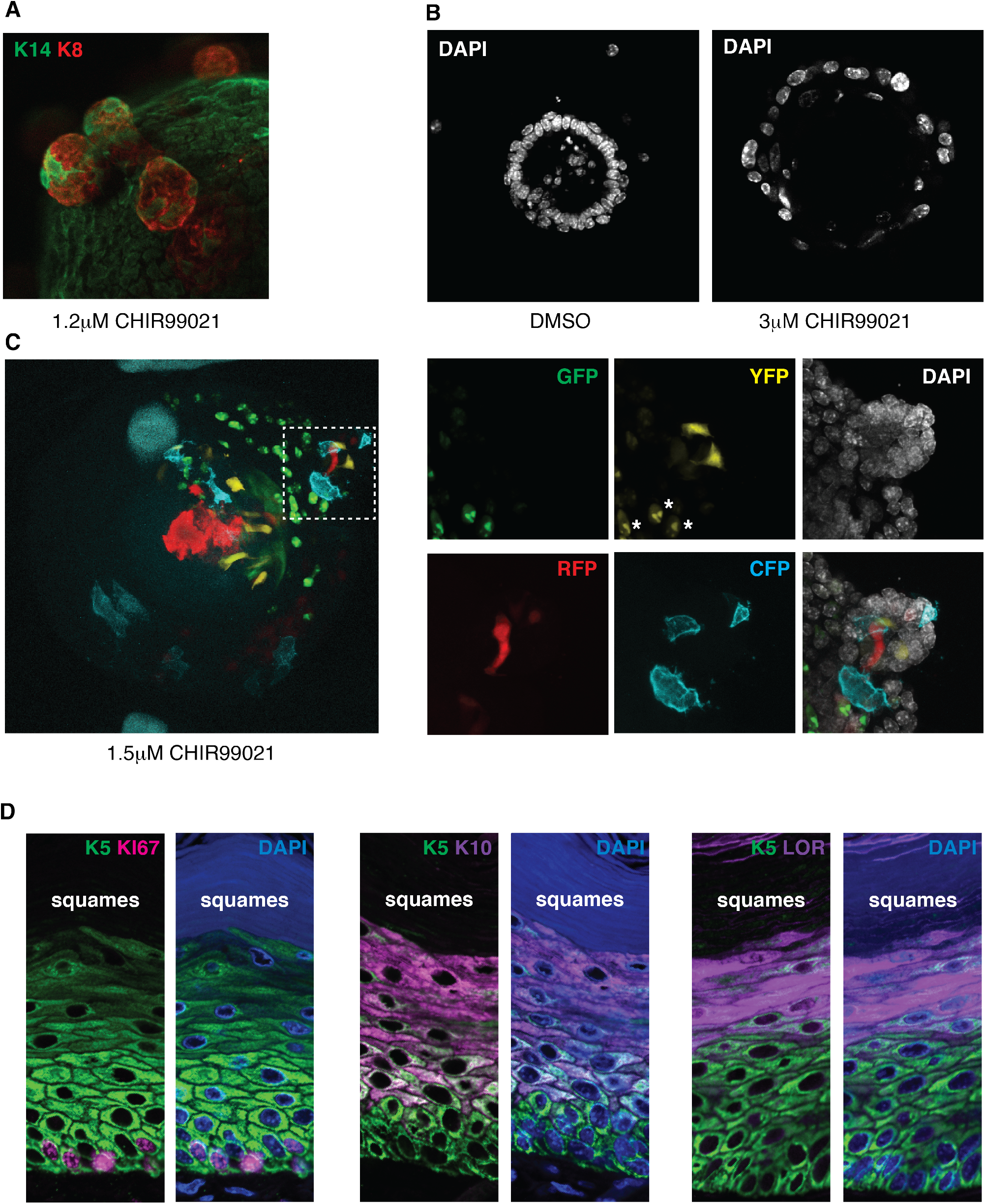
A) Wholemount confocal microscopy image showing protrusions of luminal cells (K8 = KRT8, luminal marker; K14 = KRT14, basal marker). B) Wholemount confocal microscopy image showing a cross section of a representative control and a representative WNT_high_ organoid, revealing changes in nuclear shape and size. Similar changes can be observed in main Figures 1I, 2J and 3I. C) Wholemount confocal microscopy image showing lineage tracing of WNT/CTNNB1 responsive cells in a WNT_med_ *Axin2^CreERT2^;Rosa26^Confetti^* organoid, revealing the outgrowth of multiple independent clones. Close ups of the area in the dashed box are shown on the right. D) Confocal microscopy images of a stratified squamous epithelium (mouse vagina), illustrating the pattern of cell division (KI67 signal in K5-positive basal cells) and differentiation (K10, suprabasal cells, and LOR, flattened and cornified cells). Squames indicates the dead, keratinized material that also shows up as the eosin stained core of the Wnt_high_ organoids in Figure 3F.

**Supplementary Figure 3.**
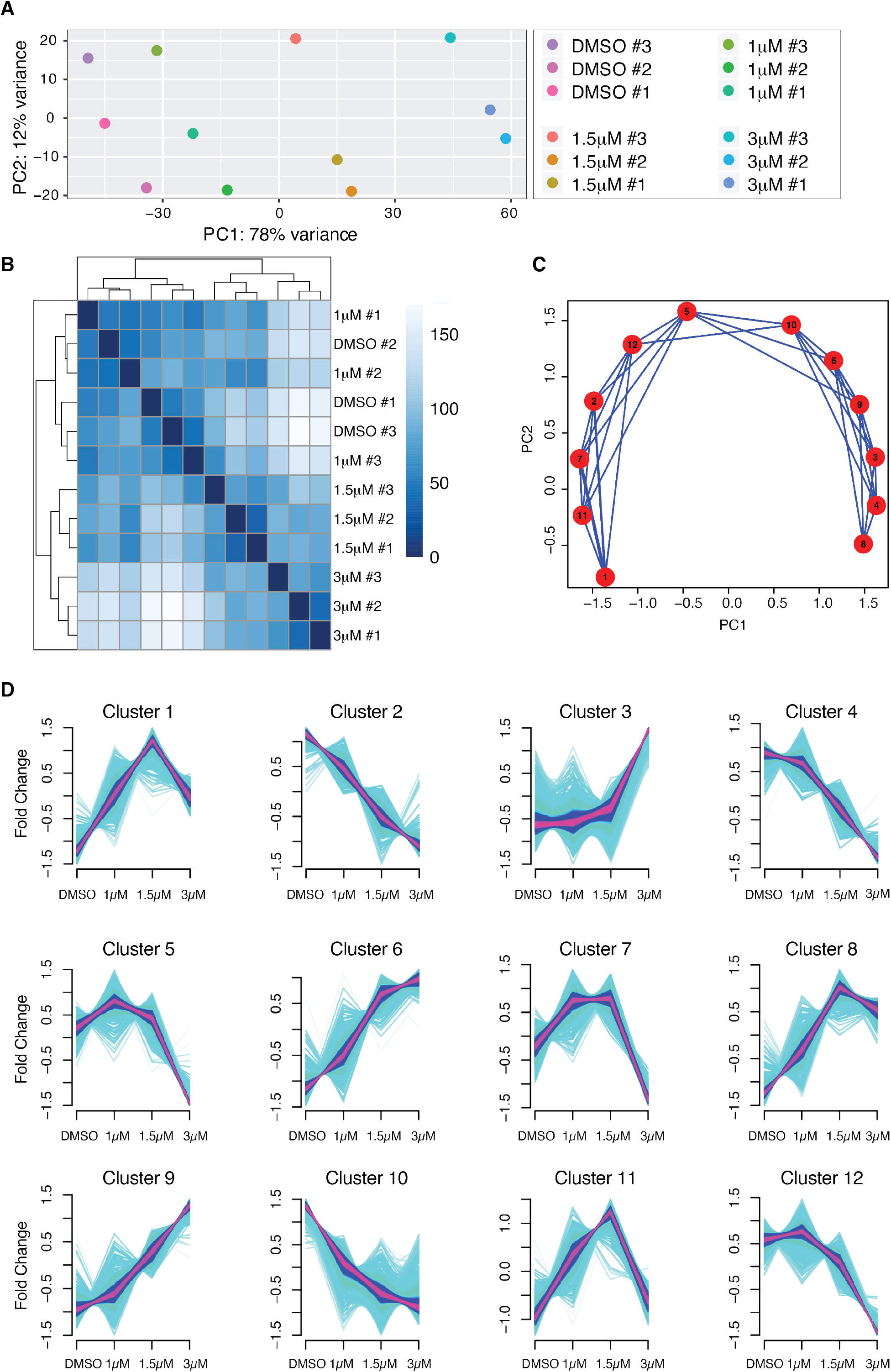
A-B) Sample level quality control of the RNAseq analysis. A) Principle component analysis (PCA) plot for the three independent RNAseq samples of the organoids depicted in Supplementary Figure 1B. B) Sample-level hierarchical clustering. C-D) mFuzz cluster analysis. C) PCA plot for the 12 different clusters. D) Gene expression changes separating the 12 different clusters. The keratinization signature was picked up in cluster 3. Loss of the mammary stem cell signature was picked up in cluster 2. This loss may seem counterintuitive, but it suggests that only low levels of WNT/CTNNB1 signaling are able to promote mammary stem-cell fate. Loss of contractile features was picked up in cluster 10.

**Supplementary Figure 4.**
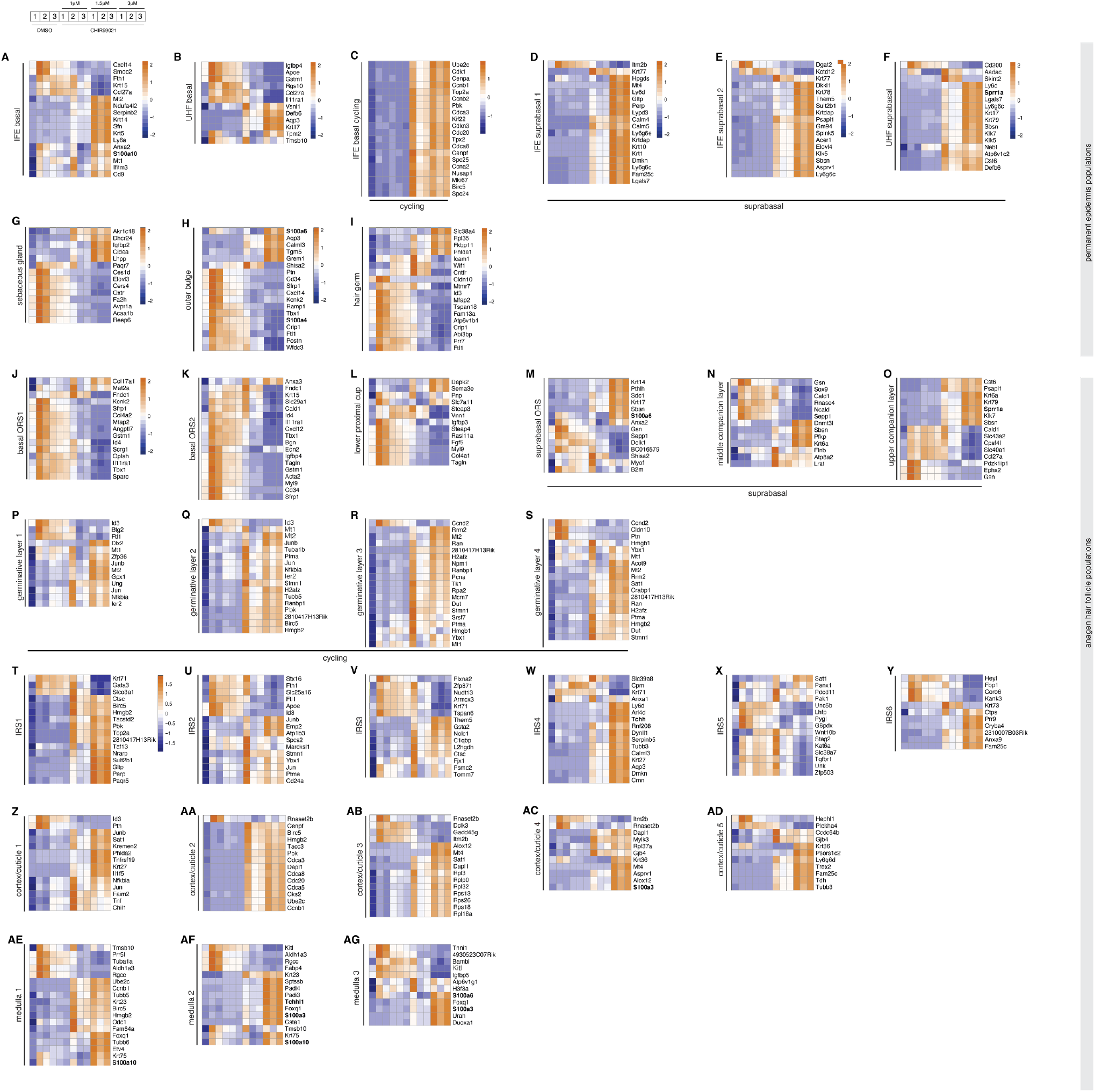
Heatmaps (unsupervised clustering, Z-score) depicting differentially expressed genes of specific cell populations of the permanent epidermis (A-I) and the anagen hair follicle (J-AG). Gene lists were generated by taking the top 20 genes for each of the populations indicated from Supplementary Table S1 of Joost et al. (2020) Cell Stem Cell. Only genes that were differentially expressed in the organoid RNAseq data are depicted. A) Interfollicular epidermis (IFE) basal, B) Upper hair follicle (UHF) basal, C) IFE basal cycling, D) IFE suprabasal 1, E) IFE suprabasal 2, F) UHF suprabasal, G) sebaceous gland, H) outer bulge, I) hair germ, J) basal outer root sheath (ORS) 1, K) basal ORS2, L) lower proximal cup, M) suprabasal ORS, N) middle companion layer, O) upper companion layer, P-S) germinative layers 1 through 4, T-Y) inner root sheath (IRS) 1 through 6, Z-AD) cortex/cuticle 1 through 5, AE-AG) medulla 1 through 3.

## MATERIALS AND METHODS

### Resource Availability

Further information and requests for resources and reagents should be directed to and will be fulfilled by the lead contact, Renée van Amerongen.

### Materials Availability

Plasmids generated in this study have been deposited to Addgene: pGlomyc-Grhl3 Addgene #172869

pGlomyc-Klf4 Addgene #172870

pGlomyc-Ovol1 Addgene #172871

### Data and code availability

The RNAseq data generated during this study are available at NCBI GEO. (GSE178321, https://www.ncbi.nlm.nih.gov/geo/query/acc.cgi?acc=GSE178321)

All R packages used for bioinformatics analysis, data plotting and summary statistics are listed in the materials and methods additional resources section.

All primer sequences are listed in the materials and methods additional resources section. Source data for all graphs and heatmaps are available in the supplementary files.

### EXPERIMENTAL MODEL AND SUBJECT DETAILS

#### Animals

All mice used for this study were maintained under standard housing conditions. Animals were housed in open or IVC cages on a 12h light/dark cycle and received food and water *ad libitum*. All experiments were performed in accordance with institutional and national guidelines and regulations and approved by the Animal Welfare Committees of the University of Amsterdam and The Netherlands Cancer Institute. All primary organoid cultures were established from FVB/NHan®Hsd mice (purchased from Envigo), except for organoids used for lineage tracing, which were established from compound *Axin2^CreERT2^;Rosa26^Confetti^* animals (own colony, mixed background).

*Axin2^CreERT2^* (Van Amerongen et al., 2012) and *Rosa26^Confetti^* (Snippert et al., 2010) strains can be obtained from Jackson labs (#018867: (B6.129(Cg)-Axin2tm1(cre/ERT2)Rnu/J; #017492: B6.129P2-Gt(ROSA)26Sortm1(CAG-Brainbow2.1)Cle/J).

#### Primary mouse mammary organoid cultures

Mammary glands (third thoracic and fourth inguinal) were harvested from 8-11 week-old virgin female mice. Mammary organoids cultures were established according to published protocols (Ewald et al., 2008; Nguyen-Ngoc et al., 2015). Briefly, fat pads were minced with scissors (∼ 20 times) and transferred to a tube with 10 ml collagenase/trypsin solution consisting of DMEM/F12/Glutamax (Gibco) supplemented with 0.02 g trypsin (Gibco), 0.02 g collagenase type IV (Sigma-Aldrich C5138), 5 ml Fetal Bovine Serum (Gibco), 250 μl of 1 μg/ml insulin (Sigma-Aldrich I6634) and 50 μl of 50 μg/ml gentamicin (Sigma-Aldrich G1397), and were incubated for 30 min at 37°C shaking at 200 rpm. The resulting suspension was centrifuged at 1500 rpm for 10 min, and then resuspended in 4 ml DMEM/F12 + 80 μl DNase (1U/μl) (Promega M6101). The DNase solution was gently shaken by hand for 2–5 min, followed by centrifugation at 1500 rpm for 10 min. Four differential centrifugations (pulse to 1500 rpm in 10 ml DMEM/F12) were performed to separate single cells (including fibroblasts) from organoids. Isolated organoids were mixed with 50 μl of Growth Factor Reduced Matrigel (Corning), seeded in an 8 well chamber slide pre-coated with 20 μl of Matrigel per well, and incubated for 30 min at 37°C. After matrigel polymerization, basic organoid growth media was added (DMEM/F12, 1% v/v insulin, transferrin, selenium (Gibco 41400045) and 1% v/v penicillin/streptomycin (Gibco, 100X stock). Organoids were cultured for 7 days in basic organoid medium and treated with either 3 μM, 1.5 μM or 1 μM CHIR99021 (BioVision/ITK Diagnostics; diluted in basic organoid medium from 6mM stock resuspended in DMSO at 1:2000, 1:4000 and 1:6000, respectively) or DMSO (VWR, 1:1000) as vehicle-control. Media was refreshed every 2-3 days and cultures were analyzed after 7 days.

#### Cell lines

The BC44 cell line (Deugnier et al., 1999) was a gift from Marie-Ange Deugnier (Institute Curie, Paris, France). Cells were cultured in RPMI medium with L-Glutamine, 10% FBS and 5 µg/ml insulin and grown at 37ᵒC and 5% CO2. For passaging, cells were trypsinized with 0.05% trypsin-EDTA for 5 minutes at 37°C, resuspended in medium and centrifuged for 4 minutes at 1500 rpm. The pellet was resuspended in fresh medium and cells were counted using a Neubauer counting chamber. Cells were passaged 1:6 or 1:10 two to three times per week.

### METHOD DETAILS

#### Organoid embedding and sectioning

Matrigel drops containing organoids were scooped from individual wells and incubated for 1.5 hours in 800 μl Cell Recovery Solution (Corning) on ice. Samples were spun down for 4 minutes at 300 rcf at 4°C. After removal of supernatant, 500 μl 4% PFA was added to each sample and incubated for 1 hour. Fixed organoids were spun down for 4 minutes at 300 rcf at 4°C and washed with Milli-Q water. The organoid pellet was then embedded in 2% agarose (Sigma) followed by an overnight incubation in 70% Ethanol. Agarose blocks containing organoids were sequentially incubated in 100% EtOH, 100% isopropanol and terpene (Histoclear, National Diagnostics), each step at room temperature for 2 hours, followed by incubation in liquid paraffin (Paraplast X-tra, Carl Roth, melting point: 50-54 °C) at 55°C. Samples were embedded in paraffin and sectioned at 5 µm.

#### Immunofluorescence

Paraffin sections were incubated for 6 minutes in terpene (Histoclear, National Diagnostics) and rehydrated in 100% isopropanol, following ethanol gradient baths (100%, 70% and 50%) and demi water. For antigen retrieval, sections were heated for 2.5 hours at 85°C with 10mM sodium citrate (pH 6.0) solution. Sections were cooled to room temperature and incubated for 30 min in 0.3% H_2_O_2_ to block endogenous peroxidase activity. After a PBS wash, sections were blocked for 1 hour with 2.5% BSA and incubated with primary antibodies overnight. The following primary antibodies were used: rat-α-K8 (1:250; TROMA-I; DSHB), rabbit-α-K14 (1:1000; PRB-155P; Biolegend), chicken-α-K5 (1:1000; 905901; Biolegend), rabbit-α-K10 (1:1000; 905404; Biolegend), rabbit-α-Loricrin (1:500; 905104; Biolegend), rabbit-α-KI67 (1:100; ab16667; Abcam). Secondary antibodies were incubated in PBS for 1 hour at room temperature. The following secondary antibodies were used: goat α-rat Alexa Fluor 647 (1:1000; A21247; Invitrogen), goat α-rabbit Alexa Fluor 568 (1:1000; A11011; Invitrogen), goat α-chicken Alexa Fluor 488 (1:1000; A11039; Invitrogen). Nuclei (DNA) were stained with 6-diamidino-2-phenylindole dihydrochloride (DAPI, Invitrogen). Slides were embedded with Mowiol (0.33 g/ml glycerol (Sigma-Aldrich 15523-1L-R), 0.13 g/ml Mowiol 4–88 (Sigma-Aldrich 81381-50G), 0.13M Tris.HCL (pH 8.5)).

#### H&E staining

Paraffin-embedded sections were incubated for 60 min at 55 °C and de-paraffinized for 6 minutes in terpene (Histoclear, National Diagnostics) and rehydrated in 100% isopropanol, following ethanol gradient baths (100%, 70% and 50%) and demi water. Slides were stained with hematoxylin (Merck Millipore) for 20-30 sec and then washed for 5 min in running tap water, following 3 min in PBS and 5 minutes in 70% Ethanol. Slides were stained with eosin (Sigma) for 3 min, washed twice in 70% EtOH for 4 min, dehydrated in 70% EtOH, 100% EtOH, 100% isopropanol and terpene and sealed with a coverslip using omnimount histological mounting medium (National Diagnostics).

#### Lineage tracing

4-hydroxytamoxifen (4-OHT, Sigma, #7904) was dissolved in 100% ethanol (1 mM stock solution). A final concentration of 1 μM 4-OHT was added to the organoid cultures on the day of plating (day 0). On the next day, the media was replaced with media containing CHIR99021. Organoids were fixed with 4% PFA for 15 minutes on day 7, washed with PBS, washed with 0.15M glycine in PBS, washed in PBS again, permeabilized with 0.5% Triton X-100 in PBS, washed in PBS, counterstained with TOPRO3 and mounted with Vectashield.

#### Microscopy

Brightfield images of mammary organoid cultures were taken on a Zeiss Axio Vert.A1 phase contrast microscope equipped with an Axiocam MRc. For imaging H&E stained slides, pictures were taken using an Axioscope A1 microscope with a Nikon Ri2 camera and NIS F freeware. For imaging immunofluorescence slides, pictures were taken using a Nikon A1 confocal microscope (20x water immersion objective with an NA of 0.75) and NIS elements AR software. Excitation with 405nm (DAPI), 440nm or 458nm (CFP) 488nm (GFP or Alexa488), 514 nm (YFP), 561 nm (RFP or Alexa561) and 633 nm (TOPRO3 or Alexa647) laser lines. For wholemount confocal imaging of fluorescently labelled organoids, images were taken on a Leica SP5 or SP8 with AOBS.

#### RNA sequencing

Matrigel drops containing organoids were scooped from individual wells of an 8-well chamber slide after 7 days of culture in the presence of either DMSO or CHIR99021 and lysed in 1 ml of Trizol Reagent (Life Technologies). RNA extraction, purification, sequencing and data processing until read count calculation were performed at the NKI Genomics Core facility as part of a collaboration with Dr. Jos Jonkers. Briefly, RNA was extracted using the Qiagen RNeasy column purification kit. RNA quality was checked with a Bioanalyer (Agilent), after which polyA+ stranded RNA library preparation was performed using the Illumina TruSeq stranded RNA prep kit. RNA-sequencing was performed on a HiSeq 2500 (Illumina) System using a stranded protocol. Single-end reads (65 bp) were aligned to reference sequence GRCm38/mm10 with Tophat version 2.1 and Bowtie version 1.1.0 (Trapnell et al., 2009). Expression values were determined by HTSeq-count (Anders et al., 2015). Original .bam files and raw counts have been deposited at NCBI GEO and are available under accession number GSE178321.

#### Bioinformatics analysis

Raw gene-level count tables were processed in R (R Core Team and Team, 2020), using DESeq2 (Love et al., 2014). No pre-filtering was performed. Normalized values were extracted from the DESeqDataSet (dds) object (provided in Supplementary File 1 as “annotated_normalizedcounts”).

Differentially expressed genes were extracted using pairwise comparisons of different treatment conditions with padj <0.1 as a cut off for false discovery (provided individually in Supplementary File 1). For Figure 2A each differentially expressed gene list was sorted according to the log2 fold change and the top 50 activated and top 50 repressed genes were selected. The resulting gene lists were combined, giving a total of 319 genes. The log2 transformed normalized expression values were used to construct the heatmap depicted in Figure 2A.

Genes meeting the criteria for differential expression in one or more comparisons (11714 genes total, provided in Supplementary File 1 as “normalized_values_DE_genes”) were used to extract gene lists and plot heatmaps using the pheatmap package (Raivo Kolde, 2019). All heatmaps were made using Z-score scaling and unsupervised clustering along rows, except for Figure 4A where genes are depicted according to their chromosomal location. With the exception of Figure 2A, dendrograms were removed in the final figures to save space.

To detect patterns of gene expression changes in our data, all genes that were differentially expressed in one or more conditions were further analyzed in R using the Mfuzz package (Kumar and Futschik, 2007). Briefly, normalized readcounts were averaged per condition (DMSO, 1 μM CHIR99021, 1.5 μM CHIR99021, 3 μM CHIR99021) and the concentrations were converted to pseudotime (0, 10, 15, 30). The fuzzifier m was estimated based on the expression set, which returned a value of 2.5. The optimal number of clusters was determined empirically and set at 12 for the final analysis. Genes making up the core of each cluster (based on a membership value >0.7) were extracted.

For gene ontology and gene set enrichment analyses, gene lists of interest (provided in Supplementary Figure 2) were analyzed using “Investigate gene sets” at http://www.gsea-msigdb.org (Liberzon et al., 2015; Subramanian et al., 2005). All 9 collections were queried: H (Hallmark gene sets), C1 (positional gene sets), C2 (curated gene sets), C3 (regulatory target gene sets), C4 (computational gene sets), C5 (ontology gene sets), C6 (oncogenic signature gene sets), C7 (immunologic signature gene sets), C8 (cell type signature gene sets). All gene set names showing specific enrichment are listed in Supplementary File 2.

To find putative regulatory transcription factors, gene sets were analyzed using Enrichr (https://maayanlab.cloud/Enrichr/) (Chen et al., 2013; Kuleshov et al., 2016). The following collections were queried: ChEA 2016, ENCODE and ChEA Consensus TFs, ARCHS4 TFs Coexp, TF Perturbations. All factors identified are listed in Supplementary File 3.

TCF/LEF ChIPseq data for mouse TCF7 (17 tracks total), TCF7L1 (1 track total), TCF7L2 (14 tracks total) and LEF1 (4 tracks total) were downloaded from https://chip-atlas.org/ and visualized in the IGV browser (https://software.broadinstitute.org/software/igv/) (Robinson et al., 2011) aligned to mm10.

#### DNA cloning

PCR based cloning was used to amplify coding regions of candidate master regulator genes (*Ovol1*, *Grhl3*, *Klf4*) from cDNA of BC44 cells treated with 3 μM CHIR99021 for 4 hours. Primers were designed with overhangs containing restriction enzyme sites of BamHI and EcoRI to enable restriction-enzyme based cloning. PCR amplification was performed using Phusion High-Fidelity DNA Polymerase (2 U/µl; Thermo Fisher, F-530L). For this 2 μl of cDNA were mixed with 10 μl of HF buffer, 5 μl dNTPs, 2 μl of the forward primer and 2 μl of the reverse primer, 0.5 μl of Phusion, 1.5 μl of DMSO and MQ sterile water up to a final volume of 50 μl. The PCR program used was the following: 95°C for 30s, followed by 34 cycles of 95°C for 10s, 55-72°C for 15s, 72°C for 60 s and a final incubation step at 72°C for 10 minutes.

The PCR product was checked on a 1% agarose gel for bands of the expected sizes (*Ovol1*= 800 bp*, Grhl3* = 1800 bp*, Klf4* = 1452 bp). PCR products were purified using a GeneJET PCR purification kit (Thermo Fisher) or a GeneJET gel extraction kit (Thermo Fisher) when purified from the agarose gel, and digested with BamHI (ER0051, Thermo Fisher) and EcoRI (ER027, Thermo Fisher) for 2h at 37°C). The pGlomyc3.1 vector (Van Amerongen et al., 2004; Jonkers et al., 1999) was digested with the same enzymes (BamHI and EcoRI) for at least 4 hours at 37°C. After digestion, Alkaline Phosphatase (AP) treatment was performed on digested vector to prevent recircularization during ligation, adding 1 μl of FastAP enzyme (Thermo Fisher, EF0654) and 2 μl of 10X FastAP buffer (c_f_ = 1X) (Thermo Fisher, #B64) and incubating it at 37°C for 10-15 min.

Digested PCR products and digested vector backbone were checked on a 1% agarose gel and purified using a GeneJET gel extraction kit (Thermo Fisher) or purified directly from the mix without running it on the gel using a GeneJET PCR purification kit (Thermo Fisher). Purified digested PCR product and digested dephosphorylated vector were ligated together for 2 hours at room temperature using a 1:1 and a 1:3 vector:insert ratio. Transformation of the ligated products was performed by mixing 5 μl of DNA with 25 μl of DH5α competent *Escherichia coli* cells. A vector only control (digested and dephosphorylated) was used as a negative control. A mixture of DNA and bacteria was incubated on ice for 15 min, heat shock was performed in a 42⁰C water bath for 1 min and the mixture was returned to ice. Afterwards, 250 μl of Lysogeny broth (LB) was added to each transformation and incubated at 37°C for 30 min. After incubation, 100 μl of the mixture was plated on Ampicillin (Amp) LB agar plates and incubated overnight at 37°C. From each condition, 12 single colonies were picked. Plasmid DNA was purified from miniprep cultures using a GeneJET plasmid DNA miniprep kit (Thermo Fisher, K0502). Constructs were checked with a control digestion performed using the same restriction enzymes used for ligation (BamHI and EcoRI). Samples containing DNA bands of the expected sizes for insert (*Ovol1*= 800 bp*, Grhl3* = 1800 bp*, Klf4* = 1452 bp) and vector (∼ 6 Kb) were sequenced verified. One miniprep of each construct cloned was selected based on sequencing results and maxipreps were prepared using a GeneJET Plasmid DNA Maxiprep kit (Thermo Fisher, K0491) to obtain a high yield of plasmid DNA. Sequencing revealed no mutations, except for pGlomyc-Klf4, where a silent mutation was found in codon 133 (CCG à CCA, proline).

#### BC44 cell treatment and transfection

For the experiments depicted in Figure 4F, cells were treated with 3 µM CHIR99021 (DMSO as a vehicle control) or 50 ng/ml purified Wnt3a (RnD) (BSA as a vehicle control) for 4 or 24 hours prior to harvesting. For the experiments depicted in Figure 4G, BC44 cells were plated in a concentration of 150,000 cells/well in 6-well plates and transfected the next day with 2 μg of the designed plasmids and X-tremeGENE HP DNA Transfection Reagent (Sigma) using a 1:1 ratio of μl X-tremeGENE HP DNA Transfection Reagent to μg DNA. First, DNA was diluted in Opti-MEM reduced serum media (GIBCO) to a final concentration of 1 μg plasmid DNA/100 μl medium (0.01 μg/μl); then X-tremeGENE HP DNA Transfection Reagent (1 μl/μg DNA) was vortexed and added without touching the walls of the Eppendorf tube; the mix was incubated for 20 minutes at room temperature. Cell culture media was refreshed before adding the transfection mix in a dropwise manner. Empty pGlomyc_3.1 vector was transfected as a negative control. Cells were harvested 24 hours after transfection.

#### cDNA synthesis and qRT-PCR

BC44 cells were lysed in 1 ml TRIzol (Invitrogen) and processed according to the manufacturer’s instructions. Briefly, the cells were lysed in the tissue culture plate for 1 minute after which the lysate was transferred to an Eppendorf tube and incubated at RT for another 5 minutes. Next, 200 µl of chloroform was added to the RNA lysates. Tubes were vortexed briefly and incubated at RT for 3 minutes. Samples were then centrifuged for 15 minutes (12,000g at 4°C). The aqueous phase was transferred to new tubes, after which 500 µl of isopropanol and 1 µl of glycogen were added. Tubes were vigorously shaken every minute for 10 minutes total, after which they were centrifuged for 10 minutes (12,000g at 4°C). The RNA pellet was washed twice by removing the supernatant, adding 1 ml of 75% ethanol, vortexing briefly and centrifuging 5 for minutes (12,000g at 4°C). After the second wash step, the ethanol was taken off and the pellet was left to air dry. The RNA was dissolved in 20 µl RNase free water and heated for 15 minutes at 55°C. RNA concentrations and purity were measured using a NanoDrop.

To normalize RNA input for qPCR analysis 2 µg of RNA was used for cDNA synthesis. Reverse transcription was performed using the SuperScript IV First-Strand Synthesis System (ThermoFisher Scientific), according to manufacturer’s protocol. Random hexamer primers were used for reverse transcription. Briefly, RNase free water was added to 2 µg of RNA until a volume of 14 µl was reached. To this mix, 4 µl of 5x SSIV buffer and 2 µl of DNase (RQ1, Promega) was added. The samples were then incubated for 30 minutes at 37°C. After this, 2 µl DNase stop solution was added and the samples were incubated again, this time for 10 minutes at 65°C. After incubation, 2 µl random hexamer primers and 2 µl dNTPs were added to each sample. Samples were then incubated for 5 minutes at 65°C. After incubation, the samples were stored on ice until cold. To each sample, 8 µl RNase free water, 4 µl 5x SSIV buffer, 2 µl DTT and 0.2 µl RiboLock enzyme was added. Samples were mixed and 20 µl was transferred to new tubes. 1 µl of SuperScript IV Reverse Transcriptase (RT) was added to the new tubes. No enzyme was added to the remaining mix (-RT control). Both the +RT and -RT mixes were incubated for 10 minutes at 23°C, 10 minutes at 55°C, and lastly 10 minutes at 80°C. The cDNA was then diluted with 180 µl RNase free water.

Per sample, a mix was made containing 10 µl RNase free water, 4 µl 5x HOT FIREPol EvaGreen qPCR Supermix (Biotium), 0.5 µl forward (10 µM stock) and 0.5 µl reverse primer (10 µM stock). This was pipetted into 96-well plates (0.2 ml), after which 5 µl cDNA was added with clean filter tips. The plate was covered with MicroAmp adhesive film and centrifuged for 5 minutes at 1500 rpm.

Quantitative PCR reactions were run on a QuantStudio 3 (Applied Biosystems) using the following program: Hold stage (2 minutes on 50°C, 15 minutes on 95°C), PCR stage (15 seconds on 95°C, 1 minute on 60°C, 40 cycles) and the Melt Curve stage (15 seconds on 95°C, 1 minute on 60°C, 1 second on 95°C). For every experiment, *Rpl13a* was used as a reference gene. Real-time PCR quantification of gene expression was performed in triplicate with one -RT control for each cDNA sample and primer pair.

To calculate relative expression of the genes of interest, the comparative quantification (ΔΔCt) method was used. To this end, the mean Ct value of the technical triplicate values was calculated for all samples. *Rpl13a* mean Ct values were used for normalization and vehicle control treated samples were used as a calibrator in each experiment. To compare different experiments, all vehicle controls were set to 1 and fold changes over vehicle control were calculated for treated samples. Individual qPCR experiments were analyzed in Microsoft Excel. Data from n=2-6 individual experiments (performed by two independent experimenters) were pooled for the final graphs in Figure 4.

#### Primer sequences

The following primers were ordered from IDT:

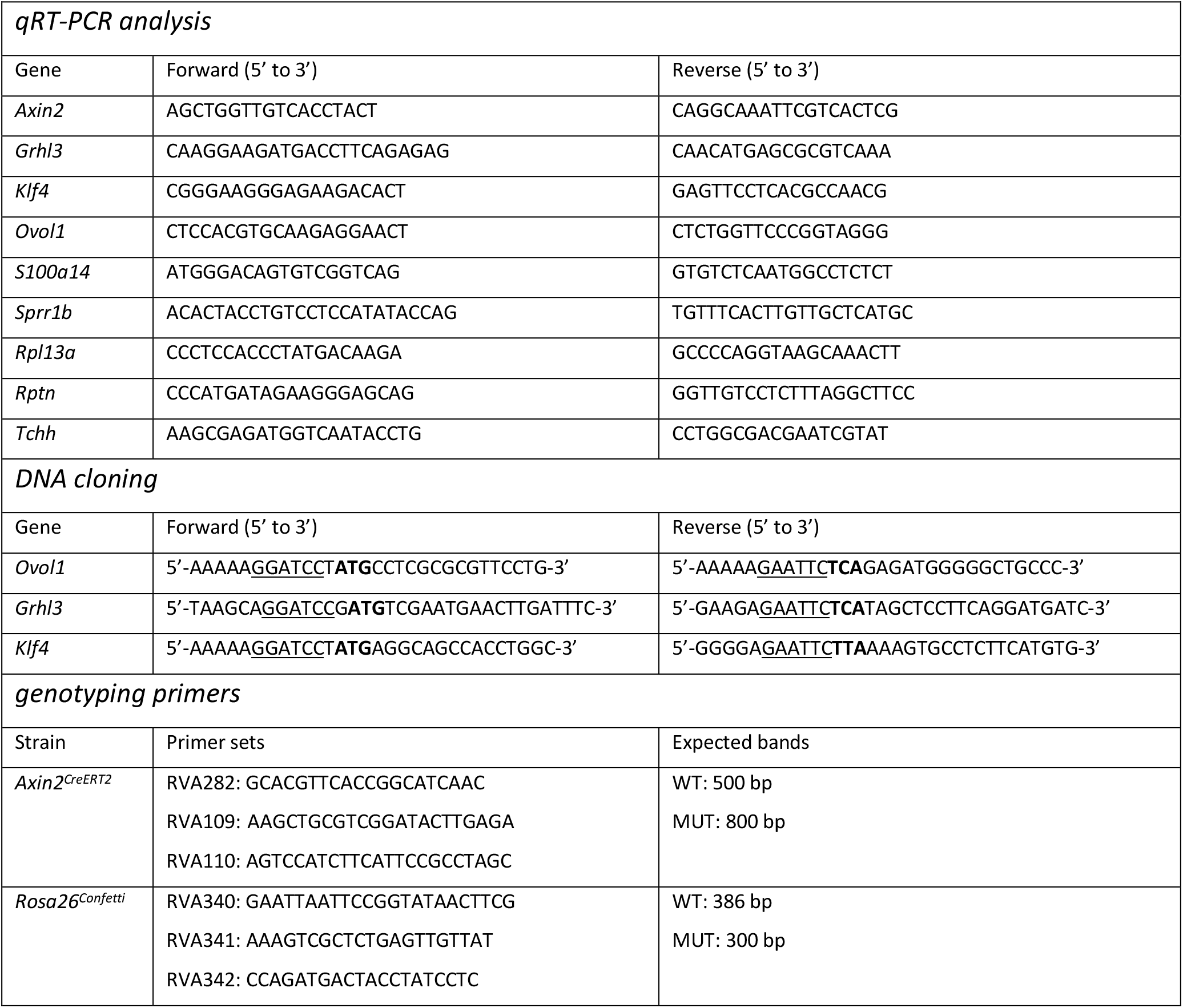

### QUANTIFICATION AND STATISTICAL ANALYSIS

All details (including exact value and definition of n, fold-changes/effects sizes, confidence intervals, etc.) can be found in the figure legends, main text or supplementary file 3.

No statistical methods were used to determine strategies for randomization or sample size estimation.

#### Organoid measurements (Figure 1 and Supplementary Figure 1)

A total of n=22 independent experiments (i.e. organoid cultures from individual mice) were analyzed. For every condition, 9-44 organoids were photographed and used to calculate area, perimeter and circularity in Fiji. For each image, the number of protrusions was counted by hand. Per condition, the 9-44 data points were averaged into a single number, resulting in n=22 data points total for each measurements. Mean or median numbers were plotted. Statistical testing was performed in R. Source data and summary statistics are provided in Supplementary File 3. Effect sizes for Figure 1G were calculated using Plots of Differences (Goedhart, 2019).

#### Organoid quantification (Figure 2)

To obtain the total cell number and K8-positive cells for each condition, the DAPI and K8 channels for each organoid image were loaded separately into CellPose (Stringer et al., 2021). Images were calibrated and segmented based on the nuclear or cytoplasm model, respectively. If necessary, the model thresholding was edited manually for each individual image to achieve optimal segmentation. The segmentation of each image was saved as a mask image that can be imported to ImageJ as ROIs via the LabelstoROIs plugin (Waisman et al., 2021). The total number of ROIs for the DAPI channel corresponds to the total number of cells present in each image, and the total number of K8 ROIs corresponds to the total number of luminal cells. The total K8 negative population, which contains both cells of a mammary and an epidermal cell fate, is obtained by subtracting the number of K8 ROIs from the DAPI ROIs. The ki67 signal was thresholded manually in FIJI and converted to a binary (dividing/non-dividing) signal. Before counting the ki67+ cells, either the DAPI or K8 ROIs were eroded by 1 pixel to exclude overlap between individual cells and to minimize the cytoplasm in K8 ROIs. The Ki67 signal in the DAPI and K8 ROIs was subsequently measured. All measurements were Excel. The total number of ROIs with a positive (255) median signal were counted to determine the number of ki67 positive cells. Images from n=2 (DMSO) or n=3 (all other conditions) independent organoid cultures were counted. Because different cell numbers were counted for each experiment and experimental condition, the data were averaged into a single data point for each individual experiment. Source data and statistics are provided in Supplementary Figure 3.

## ADDITIONAL RESOURCES

### Software

Data were handled and analyzed in Microsoft excel and R (packages: ggplot2, ggpubr, dplyr, ggbeeswarm, tidyr, rstatix, rcompanion, Rmisc, DescTools, Boot, DESeq2, Biobase, RColorBrewer, pheatmap, Mfuzz). All heatmaps and graphs were made in R Studio or Prism, except for Figure 1G which was made using the online Plots of Differences tool. Microscopy images were analyzed using Fiji and multi-channel overlays were made using the Image 5D plugin. All final figures were composed in Adobe Illustrator.

### Databases and online tools

Plots of Differences: https://huygens.science.uva.nl/PlotsOfDifferences/

CellPose: http://www.cellpose.org

MSigDB: https://www.gsea-msigdb.org/gsea/msigdb/

Enrichr: https://maayanlab.cloud/Enrichr/

ChIP atlas: http://chip-atlas.org

IGV browser: https://software.broadinstitute.org/software/igv/

## Acknowledgements

We thank the van Leeuwenhoek Centre for Advanced Microscopy (LCAM, Section Molecular Cytology, Swammerdam Institute for Life Sciences, University of Amsterdam) for the use of their facilities and LCAM staff for sharing their expertise and providing technical support. We thank Thijs van Boxtel, Saskia de Man, Tanne van der Wal and Maranke Koster for feedback on the manuscript.

